# Diversity and composition of gut microbiome of cervical cancer patients by 16S rRNA and whole-metagenome sequencing

**DOI:** 10.1101/2020.05.05.080002

**Authors:** Greyson Biegert, Tatiana Karpinets, Xiaogang Wu, Molly B. El Alam, Travis T. Sims, Kyoko Yoshida-Court, Erica J. Lynn, Jingyan Yue, Andrea Delgado Medrano, Joseph Petrosino, Melissa P. Mezzari, Nadim J. Ajami, Travis Solley, Mustapha Ahmed-Kaddar, Lauren Elizabeth Colbert, Ann H. Klopp

## Abstract

**Purpose:** Next generation sequencing has progressed rapidly, characterizing microbial communities beyond culture-based or biochemical techniques. 16S ribosomal RNA gene sequencing (16S) produces reliable taxonomic classifications and relative abundances, while shotgun metagenome sequencing (WMS) allows higher taxonomic and functional resolution at greater cost. The purpose of this study was to determine if 16S and WMS provide congruent information for our patient population from paired fecal microbiome samples.

**Methods:** Patients with locally advanced cervical cancers were enrolled on a prospective, observational clinical trial with a rectal swab sample collected prior to chemoradiation. Bacterial DNA was extracted from each sample and divided in two parts for 16S or WMS sequencing. We used measures of diversity richness and evenness as comparators of 16S and WMS sequencing. Relative abundances of the most common taxa were also compared between both datasets. Both techniques were tested against baseline patient demographics to assess associations identified with either or both methods.

**Results:** Comparative indices were highly congruent between 16S and WMS. The most abundant genera for 16S and WMS data did not overlap. Overlap was observed at the Phylum level, as expected. However, relative abundances correlated poorly between the two methodologies (all p>0.05). Hierarchical clustering of both sequencing analyses identified overlapping enterotypes. Both approaches were in agreement with regard to demographic variables.

**Conclusion:** Diversity, evenness and richness are comparable when using 16S and WMS techniques, however relative abundances of individual genera are not. Clinical associations with diversity and evenness metrics were similarly identified with WMS or 16S.

**Importance:** The gut microbiome plays an important role in regulating human health and disease. 16S rRNA gene sequencing (16S) and the whole-metagenome shotgun DNA sequencing (WMS) are two approaches to describe the microbial community. 16S sequencing via any amplicon sequencing-based method offers advantages over WMS in terms of precision (specific gene targeting). Additionally, 16S has historically been less costly due to the simplicity of library preparation and it does not require the same level read coverage as WMS. In this study, we performed both sequencing methods on a single rectal swab sample obtained from each cervical cancer patient prior to treatment. We showed that these two methods provide comparable information for diversity, evenness, and richness at higher taxonomic resolution, but are discrepant at a lower resolution. These methodological findings provide valuable information for the design and interpretation of future investigations of the role of the gut microbiome in cancer.

**Tweet:** (optional: 256 words, please submit a Tweet that conveys the essential message of your manuscript.) 16S may be sufficient for most initial studies of the gut microbiome in cancer patients, but WMS may be required for analysis of lower level taxonomy.

**Research support:** This research was supported in part by the Radiological Society of North America Resident/Fellow Award (to L.E.C.), the National Institutes of Health (NIH) through MD Anderson’s Cancer Center Support Grant P30CA016672, the Emerson Collective and the National Institutes of Health T32 grant #5T32 CA101642-14 (T.T.S). This study was partially funded by The University of Texas MD Anderson Cancer Center HPV-related Cancers Moonshot (L.E.C and A.K.).

## Introduction

The gut microbiome is increasingly recognized as a critical determinant of health and disease(1). The vast majority of microbiome analyses have utilized 16S rRNA gene sequencing (16S) which uses variable regions of the 16S ribosomal RNA gene to assign taxonomic classification and read abundance to calculate the relative frequency of the organisms within a sample(2). 16S is a reliable method for identifying the relative frequency of organisms but does not provide reliable functional information about the genes encoded by these organisms(3). As a consequence, whole-metagenome sequencing (WMS) data has been increasingly utilized with the goal of providing functional information about the organisms present. WMS analyzes large swaths of genomic information, which confers several advantages over 16S. Most notably, WMS allows for an increased depth and specificity of sequenced species as well as insights into gene abundance and metabolic capacity(4). Since WMS yields genomic information beyond the 16S rRNA gene, it also confers a better assessment of the true diversity of the sample. Thus, WMS can be used to provide species level resolution, as well as differences in presence of microbial genes, articulated pathways and metabolic functions(4). Yet, a limitation of shotgun sequence data is the large number of sequence reads which must be mapped to databases, which requires significant expertise to balance classification accuracy with discarded reads. Now, it is possible to analyze 16S and WMS microbiome data side-by-side to investigate bacterial communities as well as the abundance of associated genes and metabolic pathways(5–10). Still, the extent to which these two sequencing methods correlate with one another is a critical assumption, which should be explored thoroughly.

Few studies have had the opportunity to compare previously observed 16S gene associations with data from WMS on the same cohort of patients(2). By subjecting the same sample to both sequencing methods, we aim to investigate the reliability, validity and reproducibility of these different approaches. To do so we utilized baseline gut microbiome analysis from patients receiving standard chemoradiation therapy for cervical cancer in order to examine and compare 16S microbiome associations with WMS data on a variety of clinical variables. We deployed commonly used alpha diversity metrics (Inverse Simpson Diversity, Shannon Diversity, Camargo Evenness, Pielou Evenness, Observed Operational Taxonomic Units, and the Low Abundance Rarity Index) as well as abundance measures, to draw comparisons between the two datasets. Additionally, we submitted the datasets to unsupervised hierarchical clustering in order to assess if the microbiome profiles associated together in a similar manner, as would be expected from two datasets derived from a single sample source.

## Methods

### Study design and participants

We collected rectal swab samples from a cohort of 41 patients with newly diagnosed, locally advanced cervical cancer undergoing treatment at The University of Texas MD Anderson Cancer Center and Harris Health System Lyndon B. Johnson clinic. Patients with previous pelvic radiation or treatment for cervical cancer were excluded. This study was part of an IRB approved protocol (MDACC 2014-0543).

### Patient population and treatment characteristics

Patients were enrolled in an IRB-approved (2014-0543) multi-institutional prospective clinical trial at The University of Texas MD Anderson Cancer Center and the Harris Health System, Lyndon B. Johnson General Hospital Oncology Clinic. Inclusion criteria were newly diagnosed cervical cancer per the Federation of Gynecology and Obstetrics (FIGO) 2009 staging system, clinical stage IB1-IVA cancers, visible, exophytic tumor on speculum examination with planned definitive treatment of intact cervical cancer with external beam radiation therapy, cisplatin and brachytherapy. Patients with any previous pelvic radiation therapy were excluded.

Patients underwent standard-of-care pretreatment evaluation for disease staging, including tumor biopsy to confirm diagnosis; pelvic magnetic resonance imaging (MRI) and positron emission tomography/computed tomography (PET/CT); and standard laboratory evaluations, including a complete blood cell count, measurement of electrolytes, and evaluation of renal and liver function. Patients received pelvic radiation therapy to a total dose of 40−45 Gy delivered in daily fractions of 1.8 to 2 Gy over 4 to 5 weeks. Thereafter, patients received intracavitary brachytherapy with pulsed-dose-rate or high-dose-rate treatments. Patients received cisplatin (40 mg/m2 weekly) during external beam radiation therapy according to standard institutional protocol. Patients underwent repeat MRI at the completion of external beam radiation therapy or at the time of brachytherapy, as indicated by the extent of disease. Patients with no residual tumor on repeat MRI were considered to be exceptional responders while those with residual MRI tumor volumes ≤20% and >20% of initial volumes after 4 to 5 weeks after initiation of RT were considered to be standard and poor responders, respectively.

### Sample collection and sequencing

Rectal swabs were collected in clinic at the time of rectal examination prior to treatment using quick release matrix designed Isohelix swabs (Isohelix cat. SK-2S). We placed the swabs in 400 μL of Lysis buffer and stored them at -80°C within 1 hour of sample collection. One portion of each sample was sequenced using 16Sv4 rRNA sequencing targeting the v4 region with primer 515F-806R(11), while another portion was sequenced using WMS. 16S rRNA gene sequencing was performed through the Alkek Center for Metagenomics and Microbiome Research (CMMR) at Baylor College of Medicine. 16S rRNA gene sequencing methods were adapted from the methods developed for the Earth Microbiome Project(11). Briefly, bacterial genomic DNA was extracted using MO BIO PowerSoil DNA Isolation Kit (MO BIO Laboratories). The 16S rDNA V4 region was amplified by PCR and sequenced on the MiSeq platform (Illumina) using the 2×250 bp paired-end protocol yielding pair-end reads that overlap almost completely. The primers used for amplification contain adapters for MiSeq sequencing and single-end barcodes allowing pooling and direct sequencing of PCR products. Then gene sequences were clustered into OTUs at a similarity cutoff value of 97% using the UPARSE algorithm (12). To generate taxonomies, OTUs were mapped to an optimized version of the SILVA rRNA database containing the 16S v4 region and then rarefied at 6989 reads. A custom script was used to construct an OTU table from the output files generated as described above for downstream analyses. Here, OTUs were selected as a basis for further analysis because this method is currently the most common approach to 16s analysis in the clinical research setting.

For WMS data, genomic bacterial DNA (gDNA) extraction methods optimized to maximize the yield of bacterial DNA from specimens while keeping background amplification to a minimum were employed(13, 14). Metagenomic shotgun sequencing was performed on extracted total gDNA on Illumina sequencers using chemistries that yielded paired-end reads. Sequencing reads were derived from raw BCL files which were retrieved from the sequencer and called into fastqs by Casava v1.8.3 (Illumina). Then, paired-end reads (fastq format) were filtered to remove Illumina PhiX sequences and trimmed for the Illumina adapters by using bbduk in BBTools (version 38.34)(15). To remove host DNA contamination, the trimmed reads were then mapped to a human reference sequence database (hg38) by using Bowtie2 (version 2.3.5)(16). Taxonomic classification was performed through MetaPhlAn2(17). Also based on Bowtie2, we mapped the cleaned (unmapped to host genome) reads to a marker gene database (mpa_v295_CHOCOPhlAn_201901, updated 11/11/2019) to get an individual relative abundance table for each sample. Relative abundance tables for all samples were merged and converted to a biom format (version 1.0)(18), which was then imported into ATIMA (Agile Toolkit for Inclusive Microbial Analysis)(13) for statistical and diversity analysis. Additionally, we obtained the functional annotation of the microbial community by using HUMAnN2(6).

### Alpha Diversity Indices

We then analyzed data from both WMS and 16S using several alpha diversity metrics provided in the Microbiome R package(19) (R version 3.6.2), in order to assess the richness, divergence and evenness of the microbial communities within each patient sample. We calculated several index measures from observed OTU counts for 16S data and WMS data collected from MetaPhlAn2 analysis independently. The Shannon Diversity (SD)(20) and Inverse Simpson Diversity (ISD)(21) indexes provide a measure of the total amount of species within a given sample. The Camargo(22) and Pielou(23) Evenness indices are designed to calculate the proportionality of individual species within a sample population. A high degree of evenness would imply that the abundances of all individuals are roughly the same, or in equal proportions. Finally, the richness of the datasets was calculated using Observed operational taxonomic unit (OTU) counts and the Low Abundance Rarity (LAR) Index measures. The Observed OTUs index provides a count based on the presence of at least one read for a given species within a sample. The LAR index(19) instead characterizes the concentration of species which have low abundance within the sample.

### Comparative Statistical Analysis of 16S and WMS

We then paired each patient value from one dataset with its corresponding value for the same patient in the other dataset. The amount of agreement between the two datasets, in terms of alpha diversity measures, was then quantified using Spearman’s rank correlation coefficient (R or rho) with value of 1 indicating a perfect agreement, between two sets of variables.

To assess the consistency in reporting microbial abundance between 16S and WMS, we identified the most abundant taxa at the genus level for each sequencing method independently. Then, we compiled a list of organisms on either of the lists. Thus, the next set of comparisons were drawn using the total number of possible taxa identified by taxonomic name at all phylogenetic levels (with the exception of species).

We also analyzed the datasets individually while considering patient demographic and clinical characteristics. We analyzed six clinical variables to assess differences in diversity, evenness, and richness between groups. Binary classifications were analyzed using the independent t-test (Age, Smoking History, Histology) while multivariable classifications were analyzed using One-Way ANOVA (Ethnicity, Node Level, FIGO Stage). We also studied age and BMI as continuous variables in relation to Inverse Simpson Diversity and Pielou evenness for both 16S and WMS datasets. Consensus between the two datasets was defined as a p<=0.1 or >0.1. All analyses were conducted using R version 3.6.2 and Microsoft Excel (2016).

### Hierarchical Clustering

To further explore the consistency of the two datasets, specifically the sample grouping according to the putative taxa abundance profiles, we use unsupervised hierarchical clustering of each OTU table by the cluster software with default settings(24). The data used for clustering was limited by only using OTUs found in more than 14 samples. The obtained heatmaps were visualized by the Java TreeView software(25).

### Data Availability

Both 16S and WMS datasets will be available upon study completion and publication via the database of Genotypes and Phenotypes (dbGaP). Similarly, proprietary code will be available upon study completion and publication through GitHub.

## Results

### Taxonomic Composition and Abundance Using 16S and WMS

The number of putative taxa compiled in OTU tables was dramatically different between the technologies, and included 984 OTUs in the 16S OTU table yet only 451 in the WMS table. The WMS OTU table was not as sparse as the 16S table and had a different abundance distribution frequency display, which was close to normal (Figure 1, panel 1). The sparse 16S OTU table had significantly more rare low-abundance taxa; this feature is evident from the frequency distribution (Figure 1, panel 2) and is well-characterized for this type of dataset.

**Figure 1.**
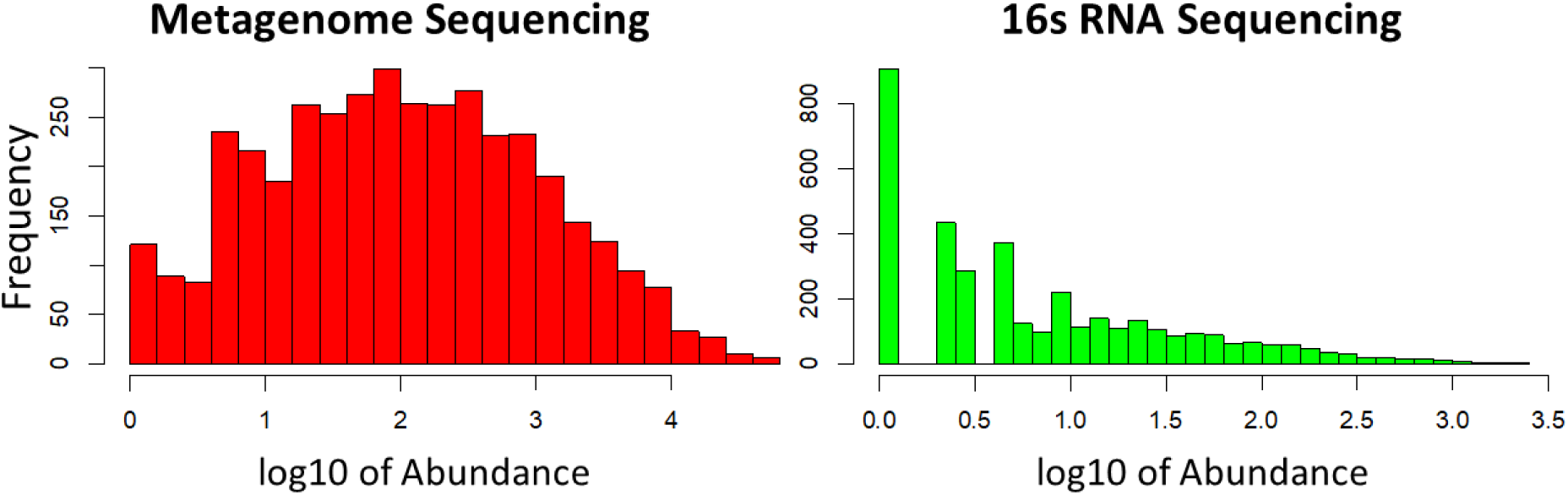
Different distribution of putative taxa species abundance in WMS and in 16S OTU tables. To display differences in abundances in terms of OTU frequencies, OTU counts were log transformed and presented here as histograms. WMS (red) OTU identifiers displayed reduced overall frequency compared to 16S (green), however the distribution of species abundance showed a more normal distribution.

The top 10 most abundant phyla and genera found in 16S and WMS are shown in Figure 2. There was better consensus between 16S and WMS datasets on the phyla level than on the genus level (Figure 2). The most abundant phyla identified in both 16S and WMS were Bacteroides, Firmicutes, Proteobacteria, Actinobacteria and Fusobacteria. Interestingly, Verrucomicrobia were found to be highly abundant only by WMS. Tenericutes were ranked third most abundant by 16S but had low abundance according to WMS. There was a significant mid-level association (rho=0.69, p=0.03) between the phyla abundances (Figure 2, panel 3). No significant associations between the abundances were identified at other taxonomic levels.

**Figure 2.**
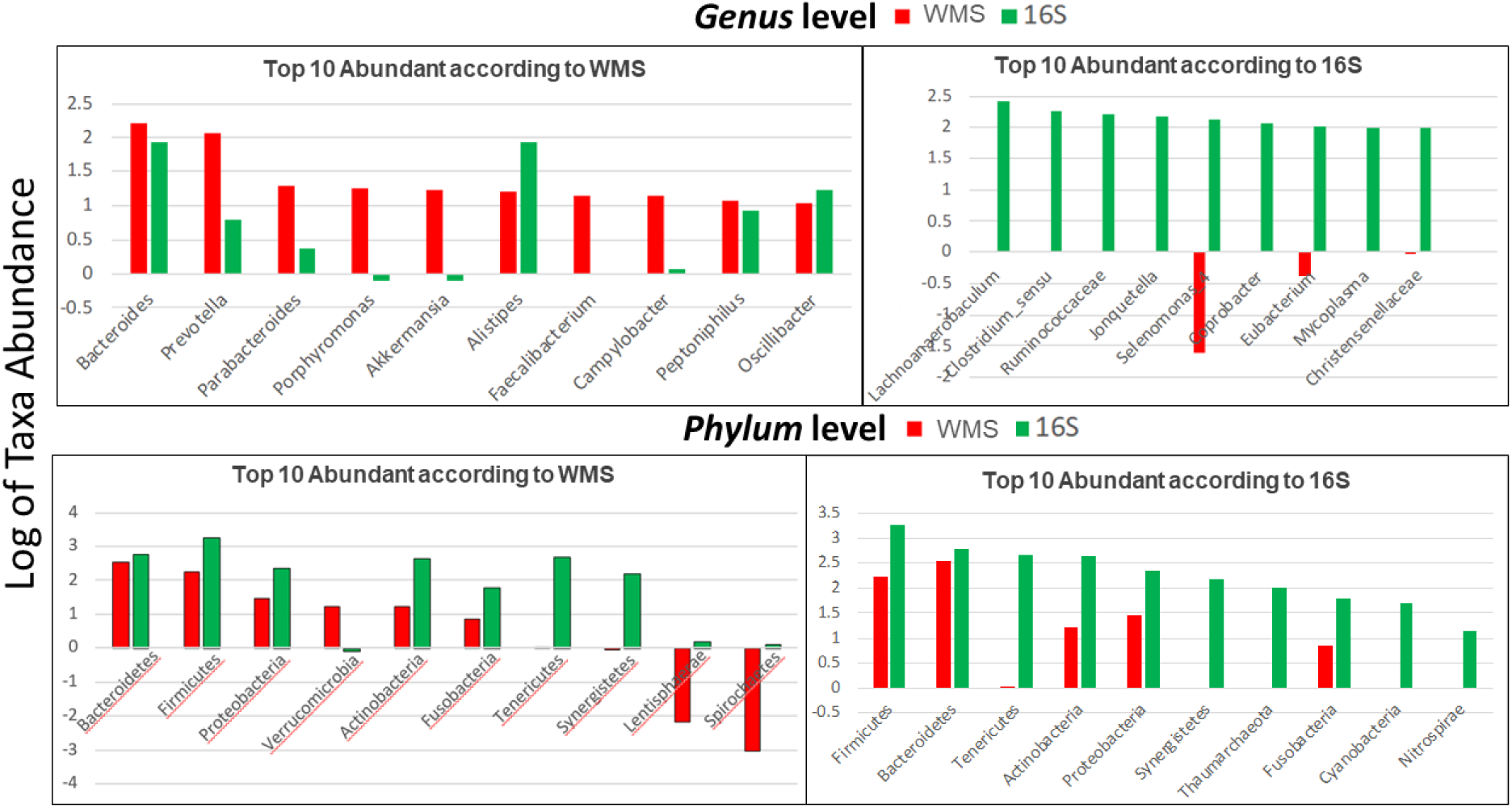
Comparison of top 10 taxa at highest and lowest taxonomic levels for WMS and 16sRNA. The bar plots presented show the top ten most abundant taxa present in the WMS (red), 16sRNA (green) as identified at the Phylum and Genus levels of taxa. The two datasets have a greater level of consensus in terms of microbial abundance at higher taxonomic levels (eg. Phylum) than lower levels (eg. Genus).

None of the top abundant genera according to 16S were identified as the most abundant genera according to WMS (Figure 2, panels 1 and 2). There was no overlap between the top 10 most abundant genera in 16S and WMS. The top 10 genera in 16S or WMS are listed in Table 1 with ranked relative abundances in each data set. Twelve genera were present in either the top 10 of 16S or WMS and present at any rank level in the other data set. Most genus level abundances correlated poorly between 16S and WMS (rho<0.15). The only genus, with relative abundance correlated well between the data sets, was Peptoniphilus (rho=0.68, p<0.01). The rest of the genera reported as the top 10 most abundant in one dataset were not present in the other dataset, and thus no abundance comparisons were made.

**Table 1.**
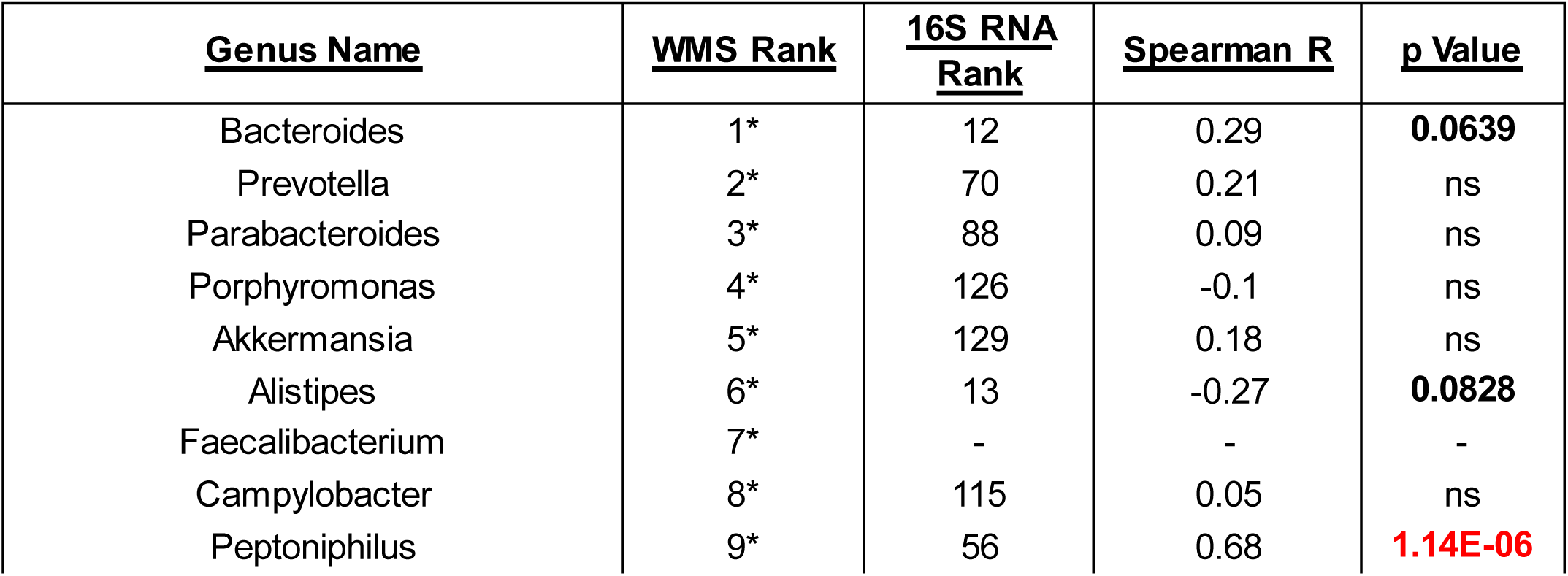
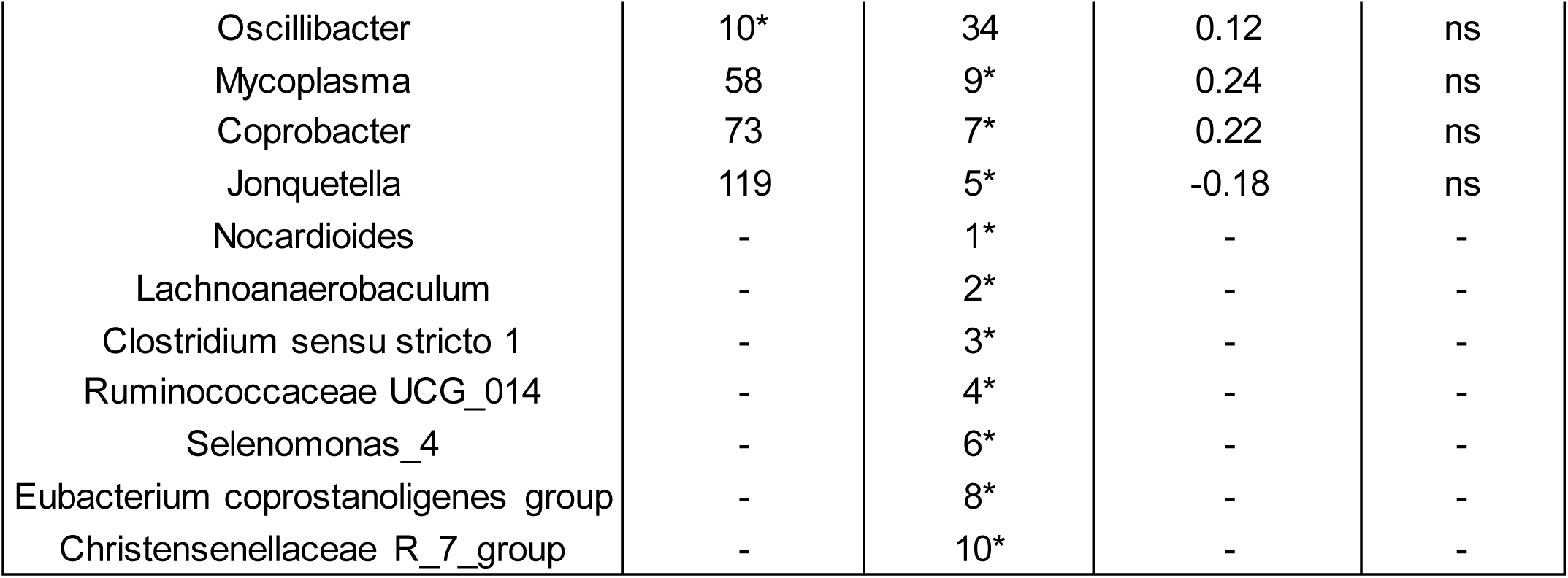
Comparisons of Top Ranked WMS and 16S Genera. The top ten most abundant Genera identified in WMS or 16S (identified with *) are shown next to their counterpart and the associated rank. Genera present in both lists were correlated in terms of abundance using Spearman R. Resulting p values are shown as not significant (ns), less than 0.1 (bold), or less than 0.05 (red). Genera present in one dataset, but not the other (-) could not be correlated.

Consistent with the difference in frequency distribution of species abundances, the 16S dataset included more putative species annotated at different taxonomic levels. Furthermore, a high percentage of the taxa identified via 16S (58-67%) were not identified by WMS, potentially as a result of using marker-gene classifiers. Conversely, most taxa found in the WMS table were also identified by 16S (Figure 3). This percentage decreased at low taxonomic levels.

**Figure 3.**
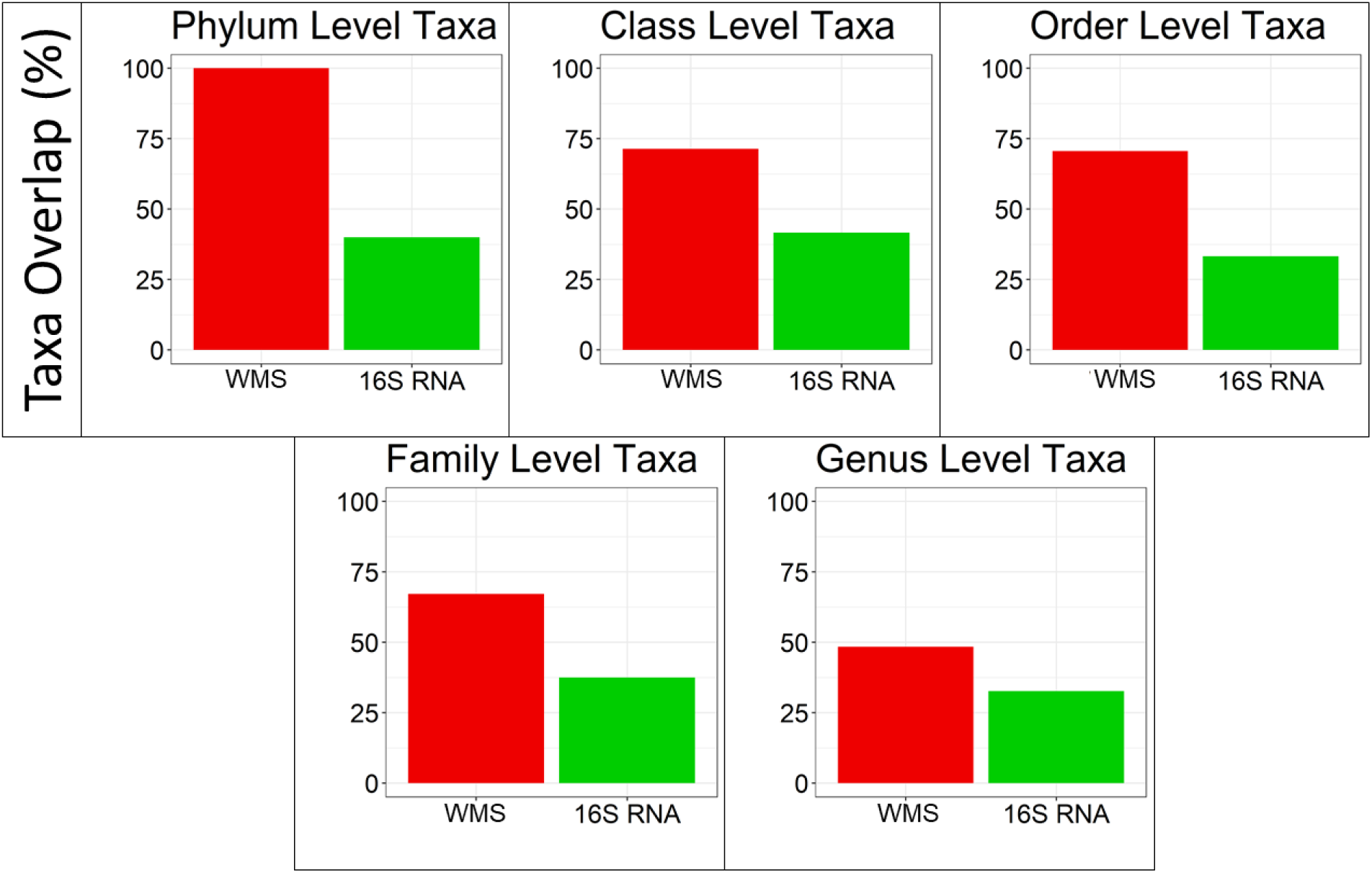
Comparison of number of overlapping taxa at each phylogenetic level for WMS and 16S RNA. The bar plots show the percentage of taxa present in WMS (red), and 16S (green) which overlap in both lists. Across all levels, many of the WMS taxa identified were also identified in the list of 16S taxa.

### Diversity, Evenness, and Richness by 16S and WMS

To further investigate the varied microbial compositions and abundances of taxa at most phylogenetic levels, we next explored the effects of different general characteristics of species diversity within the gut microbiomes. Surprisingly, we found that most indices of diversity, evenness, and richness showed significant correlation between 16S and WMS datasets (Figure 4). All of the diversity and richness measures were tightly correlated between 16S and WMS (ISD rho=0.89, p<0.01; SD rho=0.90, p=<0.01; Observed OTUs rho=0.76, p<0.01; LAR rho=0.72, p<0.001) (Figure 4). Evenness indices had a weaker correlation between 16S and WMS (Camargo rho=0.41, p<0.01; Pielou rho=0.84, p<0.01), which is not surprising considering there was greater similarity in taxa abundances in the WMS dataset then in the 16S dataset (Figure 1). Despite significant differences in the rare OTUs, low abundance rarity indexes also significantly correlated between the datasets. The slope of the regression line of the association was also consistent with a significantly greater number of rare low abundance species in the 16S dataset than in WMS.

**Figure 4.**
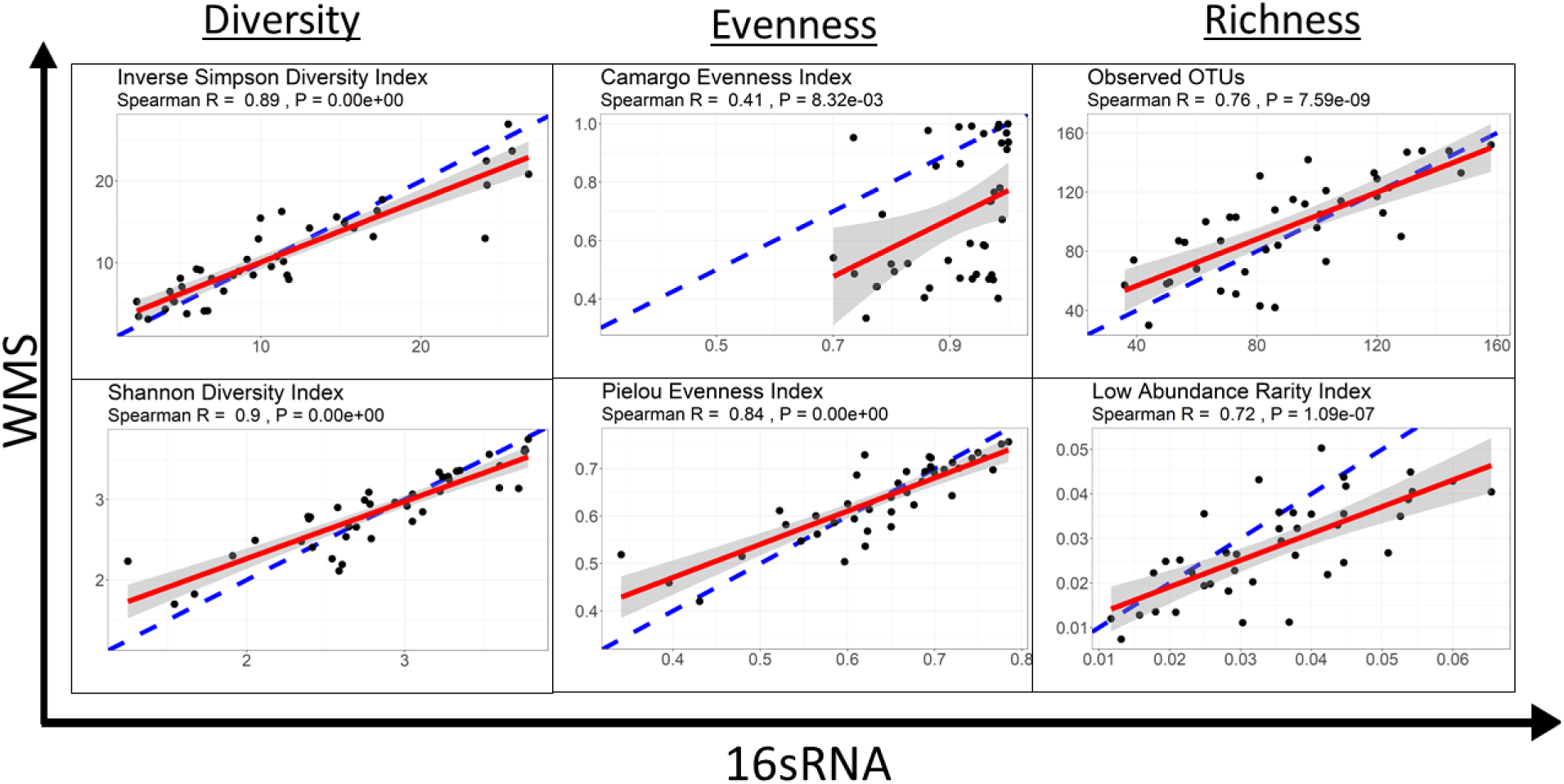
Correlation between WMS and 16sRNA in terms of diversity, evenness, and richness. In this figure, each data point represents a single patient. Consensus between both sequencing methods in terms of alpha diversity is calculated by a Spearman Correlation (R). The slope of the correlation is represented by a red line, while the 95% confidence interval is represented by a grey shaded area. The data derived from 16S sequencing correlates well with the diversity assessment values derived from WMS for diversity and richness. The evenness measures suggest that the sequencing methods differ in terms of the proportionality of individual bacterial taxa.

### Association of Demographic Characteristics with diversity of microbiomes and specific taxa

In our next step, we investigated whether the differences and similarities considered above affected biological conclusions drawn from each dataset. Namely, we explored demographic variables (Supplementary Table 1) in association with gut microbiome diversity and specific taxa using either the 16S or WMS dataset. When diversity, evenness and richness indices were compared to baseline characteristics using both 16S and WMS (Table 2), only age was associated with ISD in both WMS and 16S (p<0.1). Age was associated with SD diversity (p=0.04), and Pielou evenness (p=0.01) using 16S, but not WMS. Camargo evenness was associated with age only using WMS (p=0.008). LAR richness was associated with BMI using WMS (p=0.05) but not 16S. Other baseline demographic variables were not associated with diversity, evenness or richness using any metric. Overall, there was consensus between methods (both p<=0.1 or >0.1) across all demographics for ISD only. A positive correlation between age and gut diversity, and between age and evenness was identified in both 16S (ISD; rho=0.37, p=0.02. Pielou; rho=0.39, p=0.01) and WMS (ISD; rho=0.29, p=0.06. Pielou; rho=0.28, p=0.08) data (Figure 5). Both datasets failed to find a difference between patient populations regarding ethnicity, smoking status, tumor histology, nodal involvement, or FIGO Stage.

**Table 2.** Correlation between WMS and 16sRNA in terms of Diversity, Evenness, and Richness. In this table, patient demographics and clinical assessments were collected and used as classification criteria to investigate differences between these characteristics in terms of alpha diversity measurements discussed earlier. Both datasets were analyzed using either a parametric t-test [Smoking History (Yes/No), Histology (Adenocarcinoma/Squamous Cell Carcinoma)], linear regression [Age and BMI], or One-Way ANOVA [Ethnicity (White, Black, Hispanic, Other), Node Level (Common Iliac/External Iliac/Internal Iliac/None/Para-Aortic), FIGO Stage (IA1, IB1, IB2, IBI, IIA, IIB, IIIB, IVA)]. The resulting p value measures are indicated on the table as being either non-significant (ns), less than 0.1 (black) or less than 0.05 (red). Consensus between both methods, whole-metagenome sequencing and 16sRNA sequencing, indicates the validity of using either method for exploring that clinical variable.

| Clinical Variable | Diversity |  |  |  | Evenness |  |  |  | Richness |  |  |  |
| --- | --- | --- | --- | --- | --- | --- | --- | --- | --- | --- | --- | --- |
|  | Inverse Simpson |  | Shannon |  | Camargo |  | Pielou |  | Observed OTUs |  | LAR |  |
|  | WMS | 16sRNA | WMS | 16sRNA | WMS | 16sRNA | WMS | 16sRNA | WMS | 16sRNA | WMS | 16sRNA |
| Age | 0.0631 | 0.0166 | ns | 0.0396 | 0.0077 | ns | ns | 0.0111 | ns | ns | ns | ns |
| Ethnicity (W,B,H,O) | ns | ns | ns | ns | ns | ns | ns | ns | ns | ns | ns | ns |
| Smoking History (Y/N) | ns | ns | ns | ns | ns | ns | ns | ns | ns | ns | ns | ns |
| Histology (Adeno/Squam) | ns | ns | ns | ns | ns | ns | ns | ns | ns | ns | ns | ns |
| Node level | ns | ns | ns | ns | ns | ns | ns | ns | 0.0748 | ns | ns | ns |
| FIGO Stage | ns | ns | ns | ns | ns | ns | ns | ns | ns | ns | ns | ns |
| BMI | ns | ns | ns | ns | ns | ns | ns | ns | ns | ns | 0.0468 | ns |

**Figure 5.**
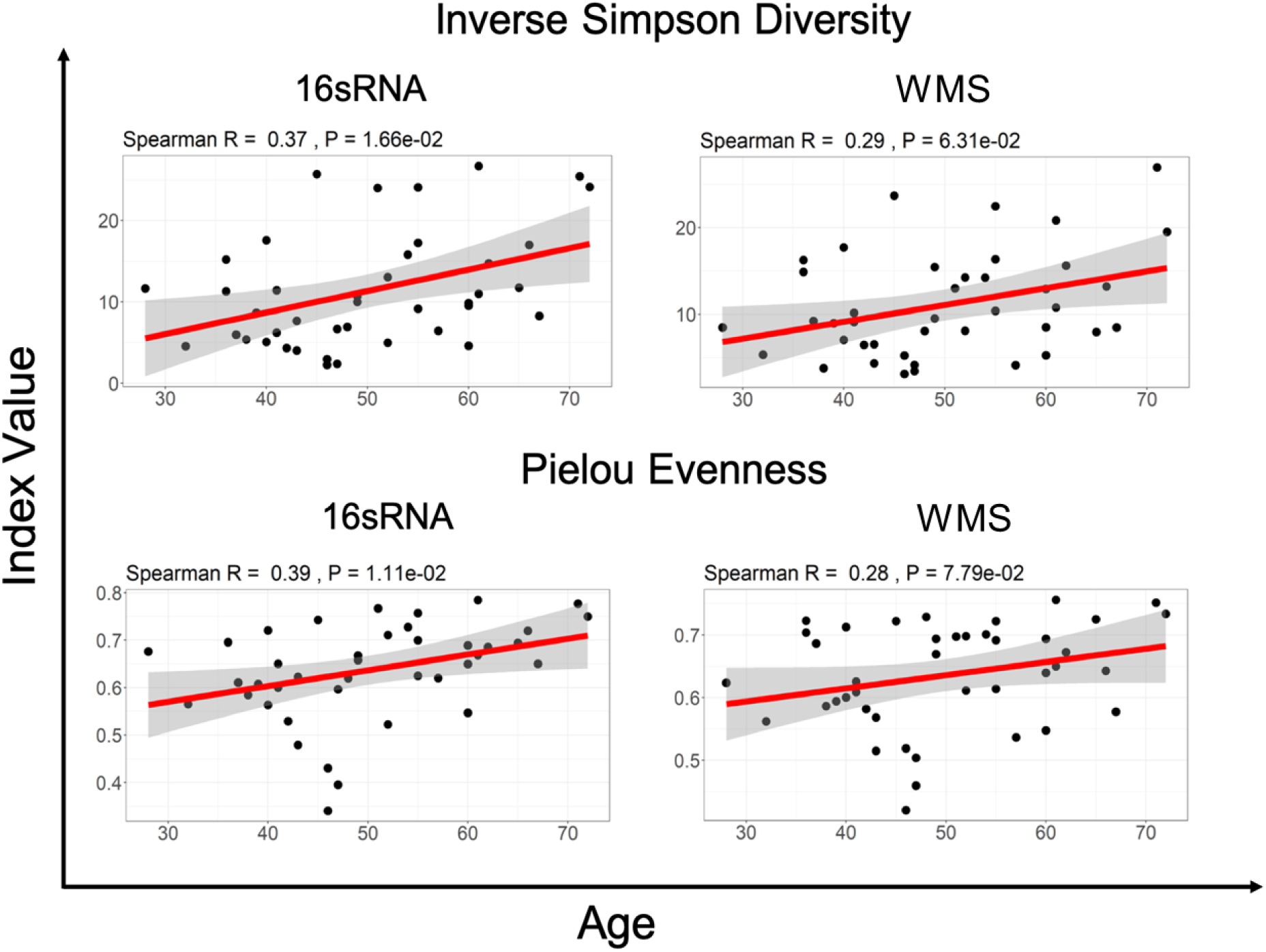
Correlation between age and Inverse Simpson Diversity and Pielou Evenness for 16S vs WMS. The slope of the correlation is shown in red while the 95% confidence interval is indicted by the grey shaded region. Spearman Correlation shows a weak association between age and the Inverse Simpson Diversity Index value as well as the Pielou evenness Index value, for both 16sRNA and WMS data.

We further explored specific taxa associated with the age of cervical cancer patients using Linear Discriminant Analysis (LDA) Effect Size (LEfSe). The clinical variable of age, was classified in three different ways: over vs under 50 years of age, over vs under the median age (49 years), and finally the patients were split into three sections where the 14 youngest and 14 oldest patients were compared against each other, with middle age group omitted (SFig. 1). We applied the one-against-all strategy with a threshold of 3 on the logarithmic LDA score for discriminative features and α of 0.05 for factorial Kruskal-Wallis test among classes. Regardless of the classification method used, the taxa identified as significantly enriched in older or younger patients was not consistent between 16S and WMS datasets.

### Grouping of cervical cancer patients in terms of putative species abundances

Unsupervised hierarchical clustering of samples based on the species abundances in WMS and 16S (Figure 6) and OTU tables revealed 2 broad groups of patients in each hierarchy with significant overlap among patients comprising each group (Fisher’s Exact Test p-value is 0.004). Despite the significant differences in the number of genera identified by 16S and WMS, the hierarchical clustering of OTUs was consistent between datasets and revealed a set of OTUs enriched with *Prevotella, Peptoniphilus,* and *Porphyromonas*. These genera were more abundant in both 16S and WMS Cluster 1, but less abundant in 16S and WMS Cluster 2. Most notably, the grouping of patients in Cluster 1 and 2 was associated with the BMI index of the patients (Fisher’s Exact Test p-value is 0.002 for WMS and 0.06 for 16S). There were significantly more patients with BMI<median (28.63) in Cluster 1 (WMS and 16S) than in Cluster 2 (both datasets). There were 20 patients in the 16S Cluster 1 and 24 patients in the WMS Cluster 1. Thirteen of which were in common between those clusters, and they were grouped close to one another. Indicating a greater degree of similarity in terms of their OTU abundance profiles.

**Figure 6.**
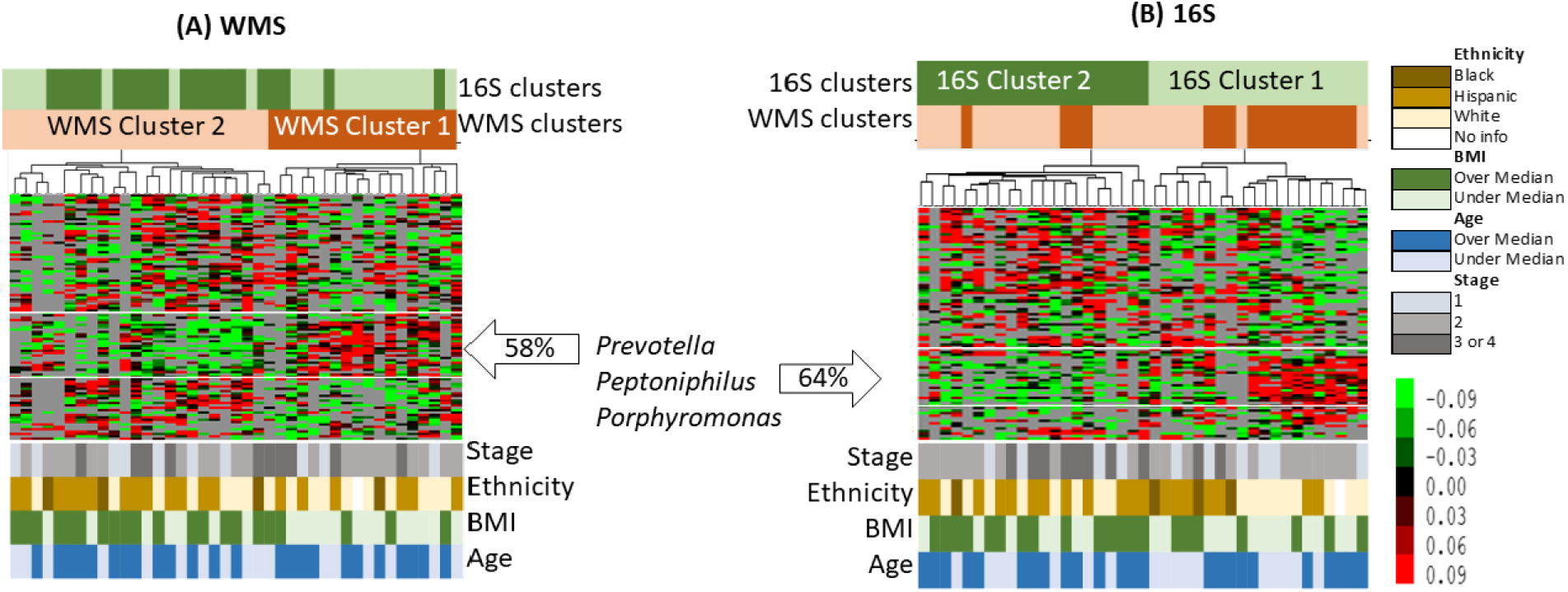
Unsupervised hierarchical clustering of samples in terms of putative species abundances identified by WMS and by 16S. To generate these heat maps, only OTUs found in more than 14 samples were considered; 91 OTUs in 16S OTU table and 103 in the WMS OUT table. Overlay between 16S and WMS sample Clusters are shown at the bars above the heat map, while demographic data is presented at the bottom.

## Discussion

This study is limited by our analytic pipelines and available samples, but the results suggest that 16S with OTU clustering provides a similar description of sample diversity and composition for gut microbiomes of cervical cancer patients versus WMS. This finding is important, as it allows researchers to analyze a larger number of samples using 16S at a fraction of the cost of WMS. Camargo evenness and skewness were the least correlated indices between the two methodologies, which suggests that the sequencing methods differ in terms of the proportionality of individual bacterial taxa. This might be improved using a 16S analysis pipeline that uses amplicon sequence variants, such as QIIME2, to retain more reads. The Camargo index has low sensitivity for variation in species diversity for sample sizes <3000, while the Pielou index is a sensitive assessment index for smaller sample sizes (<1000). Thus, the Pielou evenness index is more appropriate in terms of this sample size, and correlates well between the two datasets(20). With regards to rare taxa (LAR), WMS provides more noise in a dataset by identifying individual genes, which may be linked to unidentified bacterial species. 16S combined with OTU clustering can at best provide information at the genus level with a high degree of confidence and relies on 97% similarity clustering at the OTU level. This difference is to be expected, and could be exploited in specific analyses, such as searching for previously identified species or particular gene functions. It is reassuring that there was significant consensus between the methodologies on the higher order levels. Much of the focus in next generation sequencing analysis is placed on the smallest taxonomic level available (i.e. the genus or species level), but higher order taxa also provide valuable information.

Previous work has also posited a sizable amount of agreement between 16S and WMS sequencing techniques at higher orders of taxa(2), consistent with these results. 16S and WMS have a significant degree of correlation; however, most of those studies utilize data derived from samples collected in similar but not identical contexts. This project provides a unique opportunity in that both 16S and WMS sequencing datasets were derived from a single sample collected from each patient and then bacterial DNA was extracted for both methods. Using this high-quality information, we investigated the correlation of these two datasets in terms of microbial composition abundance and alpha diversity, to precisely determine how well these sequencing methods corroborated. Since the two datasets are derived from the same samples, association with the clinical variables should also result in the same conclusion regardless of the sequencing method used, which was again confirmed. Age is perhaps the variable most strongly associated with microbiome diversity, which was confirmed in both datasets in our study. It is also important to note, hierarchical clustering analysis showed 9 (69%) out of 13 patients in Cluster1 were white, while only 8 (31%) of the 28 patients in the rest of the cohort were white. In addition, 12 (92%) out of the 13 patients in Cluster1 had a disease stage of 1 or 2, compared to 19 (68%) out of the 28 patients in the rest of the cohort.

An important limitation of the study that we focus solely on taxonomic characteristics of the gut community. The major advantage of WMS is that it provides an opportunity to assay functional diversity of the microbiome, a capability severely lacking in 16S data. Tools such as PICRUSt(26) can infer metabolic profiles from 16S data, but they cannot truly assemble functional pathways. Yet, the most fundamental drawback of this study is due to the limitations of analytic pipelines used in each approach and the databases available for both 16S and WMS data. Tools for analyzing 16s have been developed and successfully deployed far longer than WMS analysis software, while the WMS analysis pipelines and databases are continually being developed and shared. The differences in alignment techniques and databases would account for a lot of the variation in taxa names herein. For example, by calculating OTUs we recapitulated a popular method of alignment used in this field, but in doing so the data has been collapsed at the cost of potential diversity information. Additionally, tools used for metagenomic analysis vary based on techniques used such as distance metrics and clustering approaches(15). Here, we used OTU clustering at 97% similarity using previously described methodology from the Human Microbiome Project(13), but this data could be re-analyzed using QIIME2 and amplicon sequence variant (ASV) calling(27) and result in variations in ASV vs. OTU assignment that could affect the analysis. Amplicon sequence variant calling with DADA2 denoising(28) may be a preferable system for WMS comparisons as the pipeline is more similar to how WMS reads are treated. Another important consideration is that the MetaPhlAn2 tool inherent in the Humann2 pipeline uses a relatively small fraction of the data generated, whereas another non-marker gene based identifier such as QIIME2, Kraken 2(29) or the mothur software(30) will generate a larger and more varied, spread of results. Still, MetaPhlAn2 outperformed IGGsearch(31) which was also deployed on our WMS dataset, and it remains the most popular marker-gene based tool in the metagenome field.

Another limitation to address, for this work and many others, is establishing a confident rarefication cut off for analysis. Usually this cut off value would be validated by utilizing a mock microbial community dataset to be analyzed alongside experimental data. Here, we were unable to acquire complete mock communities as such information is privileged and difficult to attain. However, the cut off value we used was selected because it was consistently stringent across both 16S and WMS datasets while retaining as much information as possible.

All this is to say, variations in approaches to metagenome assembly pipelines similarly could affect taxonomic assignment in 16S and WMS data. It is possible that a particular sequence relevant to both datasets would be classified differently during preprocessing, highlighting the necessity of universal reference databases and sequencing alignment tools and protocol consensus.

Given this variability in sequencing and data processing pipelines, the use of multiple techniques across different types of sequencing data is an excellent way to confirm consistency in conclusions. However, limited resources (e.g. material from clinical samples, bioinformatics support, time and finances) hamper the ability for this expansive and in-depth microbiome profiling for all studies. Although WMS has been demonstrated to confer significant advantages over 16S, this work suggests there is very little additional taxonomic information identified from WMS that was not identified in 16S data. This can vary depending on the context of analysis, for example method of sample collection (i.e., whole stool vs swabs) and determining the functional components of the microbiome in question.(26, 32) It is even possible that extracting DNA from the same sample at two different times, instead of splitting a single extraction as was done here, may yield slightly different results. In our work, alpha diversity assessments such as overall diversity, evenness and richness can provide meaningful, and more important, comparable information (Figure 1) when obtained with either 16S or WMS. Furthermore, the two datasets provided a high degree of consensus when these indices were subjected to statistical analysis. This suggests that for studies where overall microbiome diversity, richness and evenness are the goals of an analysis, 16S is more than sufficient to provide this information. For basic taxonomic descriptions, there was a meaningful agreement on the phyla and higher taxa levels, suggesting that 16S is also sufficient in this setting for hypothesis-generating data. Nonetheless, these two datasets did provide some differences in taxonomic assignment, particularly on the genus level, and relative abundances of individual taxonomies. This suggests that for studies where a broader repertoire of potential species are needed, both techniques may be necessary.

In all, this evidence suggests that using 16S alone may be sufficient in the clinical cancer research setting, where available patient material, time and money can be scarce. Based on these findings, we suggest 16S for the gut microbiome of cancer patients for initial diversity, richness and evenness metrics along with higher level taxonomic classification. WMS can provide a large swath of detailed microbial information, albeit with less sensitivity than 16S, and may be ideal when additional information on genus and species level identification is needed or to confirm conclusions drawn from 16S data.

## Supplementary Information

**Supplementary Table 1.** Baseline patient demographic information. Patient demographic information was collected prior to initiation of standard treatment and reported here. These demographics were used as patient categories for downstream analysis of microbial diversity, evenness and richness. Ethnicity, History of smoking, Tumor Histology, Node Level, and FIGO Stage were categorical clinical variables while BMI was used as a continuous variable. Age was used both as a categorical variable (Over vs Under 50 years of Age; Over vs Under Median Age), and as a continuous variable.

| Variable | Category | N(%) |
| --- | --- | --- |
| Age | Under 50 | 22(53.66) |
|  | Over 50 | 19(46.34) |
| Ethnicity | B | 4(9.76) |
|  | H | 19(46.34) |
|  | O | 1(2.44) |
|  | W | 17(41.46) |
| History of smoking | Yes | 17(41.46) |
|  | No | 24(58.54) |
| Histology | Adenocarcinoma | 8(19.51) |
|  | Adenosquamous | 2(4.88) |
|  | Squamous Carcinoma | 31(75.61) |
| Node Level | Common Iliac | 6(14.63) |
|  | External Iliac | 15(36.58) |
|  | Internal Iliac | 4(9.76) |
|  | Para-Aortic | 3(7.32) |
|  | None | 13(31.71) |
| FIGO Stage | IA1 | 1(2.44) |
|  | IB1 | 1(2.44) |
|  | IB2 | 6(14.63) |
|  | IBI | 2(4.88) |
|  | IIA | 2(4.88) |
|  | IIB | 19(46.34) |
|  | IIIB | 7(17.07) |
|  | IVA | 3(7.32) |
| | Mean ± SD | Range |
| Age | 49.98 ± 10.92 | 28 – 72 |
| BMI | 28.92 ± 6.14 | 17.5 – 46.7 |

**Supplementary Figure 1.**
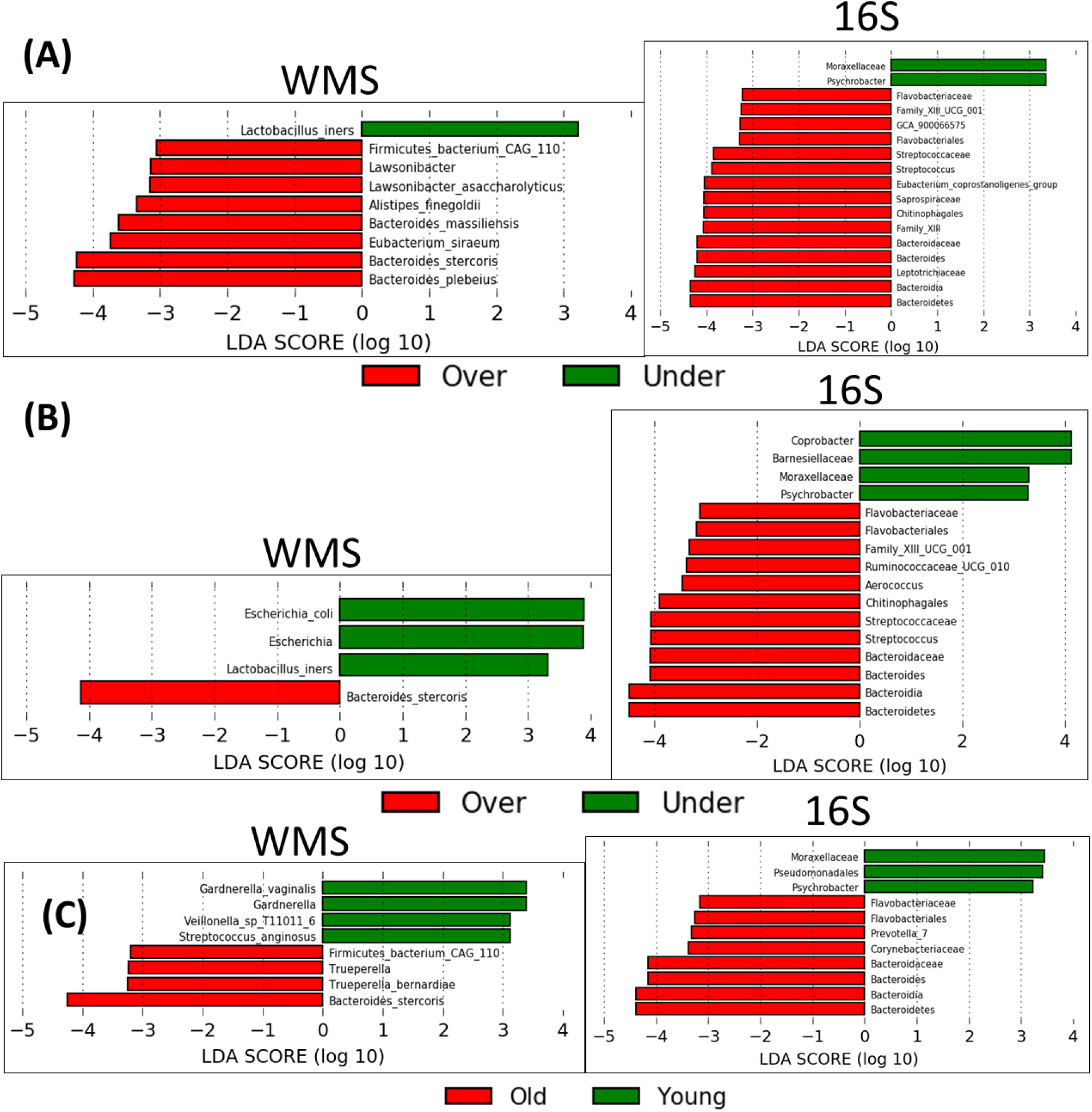
LEfSe analysis performed using 3 methods of age classification. In order to elucidate differences in enriched taxa according to the age clinical variable, patients were classified using three different metrics: A) Over vs under 50 years of age, B) Over vs under the median age of all patients, and C) the youngest third of patients (14 individuals) vs the oldest third of patients (14 individuals) in the cohort. WMS and 16S datasets were submitted to LEfSe analysis separately, and the resulting bar charts are shown here.

